# Folding Thermodynamics and Pathway Heterogeneity of Lanmodulin from Atomistic Simulations

**DOI:** 10.1101/2025.07.25.666904

**Authors:** Utkarsh Kapoor

## Abstract

Lanmodulin (LanM), a metal-binding EF-hand protein, is widely believed to fold into its functional conformation only upon coordination with rare earth elements such as lanthanide ions. Here, we challenge this assumption through advanced atomistic molecular dynamics simulations that combine parallel tempering and well-tempered metadynamics to probe the thermodynamics and folding pathways of LanM under apo conditions. By mapping the folding free energy landscape, identifying unfolded, misfolded and folded basins, and integrating contact map analysis, reactive folding trajectory analysis and diffusion map-based dimensionality reduction, we discover a striking deviation from canonical models: LanM spontaneously adopts a native-like fold, including structured EF-hand motifs and a well-formed hydrophobic core, even in the absence of metal ions. We find that the LanM folding mechanism proceeds via a two-step pathway – initial entropically driven collapse into a metastable misfolded ensemble that retain partial or full helices but in misaligned arrangements, followed by an enthalpically favorable reorganization into the folded state. Folding trajectories and diffusion map-based embeddings further reveal and distinguishes pathway multiplicity: some transitions proceed directly from the unfolded to the folded state, while others transiently occupy the misfolded basin, underscoring kinetic heterogeneity and misfolded basin’s role as an accessible but non-obligate intermediate. Taken together, these findings challenge the prevailing view of LanM as a purely metal-induced folder and instead support a model of intrinsic foldability, where native-like features emerge spontaneously and prime the protein for ion binding. Beyond refining our understanding of EF-hand protein folding, these results have direct implications for rare earth separation technologies, where the conformational readiness of apo-LanM could inspire the design of next-generation bioseparation platforms.

**Table of Contents Image:** 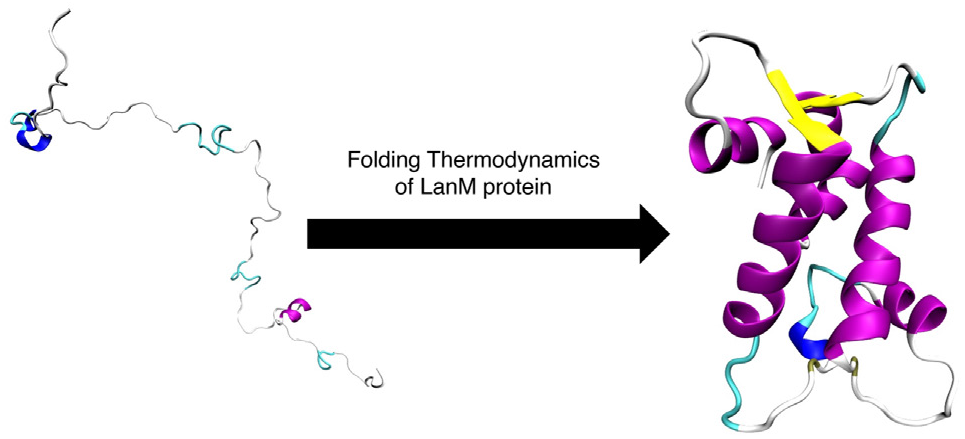

## 1. INTRODUCTION

Rare earth elements comprising the lanthanide chemical group play an irreplaceable role in modern technologies ranging from magnets and light-emitting diodes to energy-efficient batteries.[1–3] Yet their extraction and separation remain chemically challenging due to their remarkably similar ionic radii and coordination preferences.[4, 5] Recent biological discoveries have unveiled that certain methylotrophic bacteria can selectively bind and utilize lanthanide ions,[6–12] opening new avenues for biomolecular separation technologies. Among the most intriguing proteins involved is *lanmodulin* (LanM), a 12 kDa EF-hand protein discovered in *Methylorubrum extorquens* and/or *Hansschlegelia quercus*, which exhibits extraordinary affinities and selectivities for lanthanides over calcium,[6, 7] even in complex biological environments.

Previous circular dichroism, mutagenesis, and calorimetry studies demonstrate that LanM contains four EF-hand metal-binding motifs and undergoes a striking conformational transformation from a disordered to a highly helical structure, where this transition is tightly coupled to metal binding.[6, 13] Structural and biophysical studies further indicate that this folding response is not merely a passive consequence of ion binding but may underpin LanM’s function as a dynamic sensor and capture agent for lanthanides.[14] Moreover, engineered dimeric forms of LanM have been recently shown to enhance rare-earth separation efficiencies through tunable metal-induced allosteric effects.[7]

Despite detailed characterization of the holo (metal-bound) state, the thermodynamics and structural mechanisms underlying LanM folding under apo (i.e., in the absence of bound metal ions) conditions remain largely unexplored. In particular, it is unclear whether LanM remains intrinsically disordered until metal binds or whether it harbors intrinsic structural biases that predispose it toward folding even under apo conditions. Resolving this question is not only of fundamental interest in protein biophysics but is also essential to understanding how LanM achieves both high metal affinity, critical for applications in rare earth separation, where its structural preorganization could influence capture efficiency, selectivity, and kinetic responsiveness.

In this work, we use atomistic molecular dynamics simulations, enhanced with parallel tempering in the well-tempered ensemble and metadynamics, to probe the thermodynamics and folding pathways of LanM under apo conditions. By mapping the folding free energy landscape of LanM, identifying unfolded, folded and misfolded basins, and integrating contact map analysis, folding trajectory analysis and diffusion map dimensionality reduction embeddings, we provide mechanistic insights into spontaneous folding drivers of LanM and explore whether misfolded states act as kinetic traps or productive intermediates, and whether folding pathways exhibit trajectory-specific heterogeneity. Our results reveal that LanM spontaneously adopts a folded-like topology even in the absence of bound metal ions, via a two-step mechanism: an initial entropic collapse into a misfolded intermediate that retain partial or full helices but in misaligned arrangements, followed by an enthalpically driven reorganization into the folded state. Critically, this folded state exhibits the hallmarks of the holo structure, including EF-hand helices and hydrophobic core formation, even without ion coordination. We also find that folding can proceed with or without visiting the misfolded intermediate basin, suggesting that misfolded states are accessible but not obligatory. Taken together, our study challenges the prevailing view of LanM as a purely metal-induced folder and instead supports a model of intrinsic foldability with pathway multiplicity, where native-like features emerge spontaneously and enable – but are not dependent on – ion binding. These insights not only refine our understanding of EF-hand protein folding but can also inform the future design of bioinspired platforms for rare earth element recognition and separation.

The remainder of this article is organized as follows. We begin by identifying the sequence and structural determinants of LanM, followed by a thermodynamic characterization of the LanM folding landscape under apo conditions, including the delineation of unfolded, misfolded, and folded basins. Next, we analyze time-continuous folding simulation trajectories to reveal the diversity of pathways and the role of misfolded state as a metastable state but a non-obligate intermediate. We further employ diffusion map–based dimensionality reduction to uncover the low-dimensional kinetic architecture of folding transitions, enabling a more nuanced interpretation of conformational flux and basin connectivity. We conclude with a discussion of the broader structural and functional implications of our findings in the context of EF-hand metal binding proteins and rare earth element bioseparation. Lastly, we describe the computational methodologies underlying our simulations and analyses.

## 2. RESULTS

### Sequence-intrinsic determinants of folding in LanM

To contextualize the folding behavior of lanmodulin (LanM), we first examine its sequence, predicted secondary structure, amino acid composition, and charge patterning (**Figure 1**). LanM is a small ∼12 kDa protein that contains four tandem EF-hand (metal coordinating) motifs, each associated with rare earth and critical mineral binding.[6, 7] The representative structural snapshot of crystallographic reference structure (PDB: 8DQ2)[7] reveals α-helical organization, consistent with the helix-loop-helix topology typical of EF-hand domains, with short β-strands interspersed in loop regions, particularly near N- and C-terminal segments.

**Figure 1.**
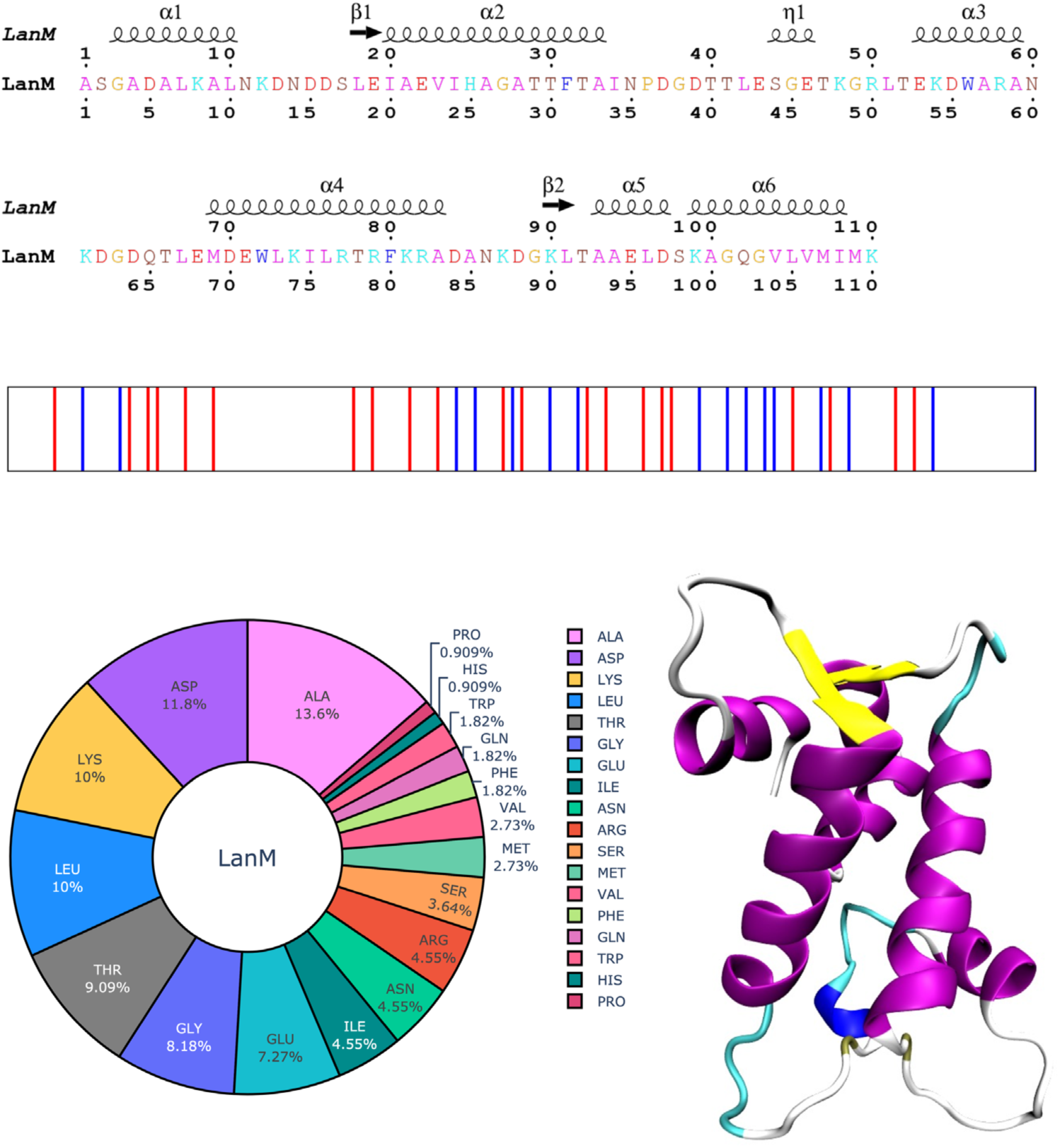
Sequence and structural features of Lanmodulin (LanM) protein. Top panel shows primary amino acid sequence and secondary structure annotation of LanM, highlighting helical segments, β-strands, and loop regions. Residues are colored by type (acidic, basic, polar, hydrophobic, etc.) for visual clarity. Middle panel shows LanM sequence charge patterning where vertical lines represent negatively charged residue (red) and positively charged residue (blue). Bottom left panel shows amino acid composition of LanM, presented as a percentage of total residues. Bottom right panel shows a representative native structure of LanM from the crystallographic model (PDB ID: 8DQ2). This structural snapshot serves as a reference to evaluate the extent to which such conformations emerge during our protein folding simulations. The sequence in the top panel was rendered using ESPript webserver[15] and the snapshot in the bottom right panel was rendered using VMD software.[16]

Previous studies by Cotruvo Jr. and co-workers have reported that LanM undergoes a large conformational change from a largely disordered state to a compact, ordered state presumably in response to presence of metal ions.[6, 7, 13, 14] However, the amino acid composition of LanM suggests that the sequence of LanM is dominated by Ala and Asp, followed by Lys, Leu, Thr, Gly, and Glu residues. The prevalence of Ala, Leu, and Thr residues intuitively suggests an inherent strong propensity for α-helix formation and hydrophobic packing. Simultaneously, the abundance of charged residues – 21 acidic and 16 basic – that leads to a net charge of –5, with a moderately negative (∼ –0.68) sequence charge decoration (SCD) parameter,[17, 18] indicates that electrostatic interactions are abundant but relatively dispersed charges rather than clustered patches. However, we note that while the acidic and basic residues are broadly interspersed, charge patterning plot also reveals short stretches of like-charged residues that alternate frequently along the sequence. Together, these characteristics indicate an inherent potential for long-range polar interactions and intra-/inter-helical contacts. Therefore, this combination of composition and sequence patterning raises a central thermodynamic question: can LanM spontaneously adopt its experimentally observed α-helical architecture without external coordination cues such as ion binding? We hypothesize that the intrinsic sequence features of LanM including its helix-forming residues and dispersed charge distribution encode sufficient structural bias to enable native-like folding even in the absence of metal ions. In this study, by simulating the folding process of LanM at atomistic resolution across a temperature range, we directly probe this hypothesis and dissect the enthalpic and entropic driving forces underlying its thermodynamic landscape.

### Thermodynamic free energy landscape of LanM

The free energy surface (FES) of LanM at 300 K, projected onto the fraction of native contacts (Q) and root-mean-square deviation (RMSD), reveals a complex yet well-defined folding landscape (**Figure 2**). Q measures the similarity between a given LanM configuration and the crystal structure and can take a value between 0 and 1, where 1 represents native-like contacts. However, one intriguing feature of the FES is the absence of conformations with Q < ∼0.45, indicating that even in the most unfolded states sampled, a non-zero fraction of native-like contacts persists. The minimum observed Q value suggests that LanM’s unfolded ensemble retains significant residual structure, likely arising from persistent short-range secondary structure elements or local helical segments that do not require long-range tertiary alignment. This is consistent with the known tendency of α-helical sequences to exhibit high local structure even in denatured conditions.[19] Since our definition of Q includes only contacts within secondary structure elements, these local short-range α-helices within flexible regions continue to contribute to Q values even in globally disordered states.

**Figure 2.**
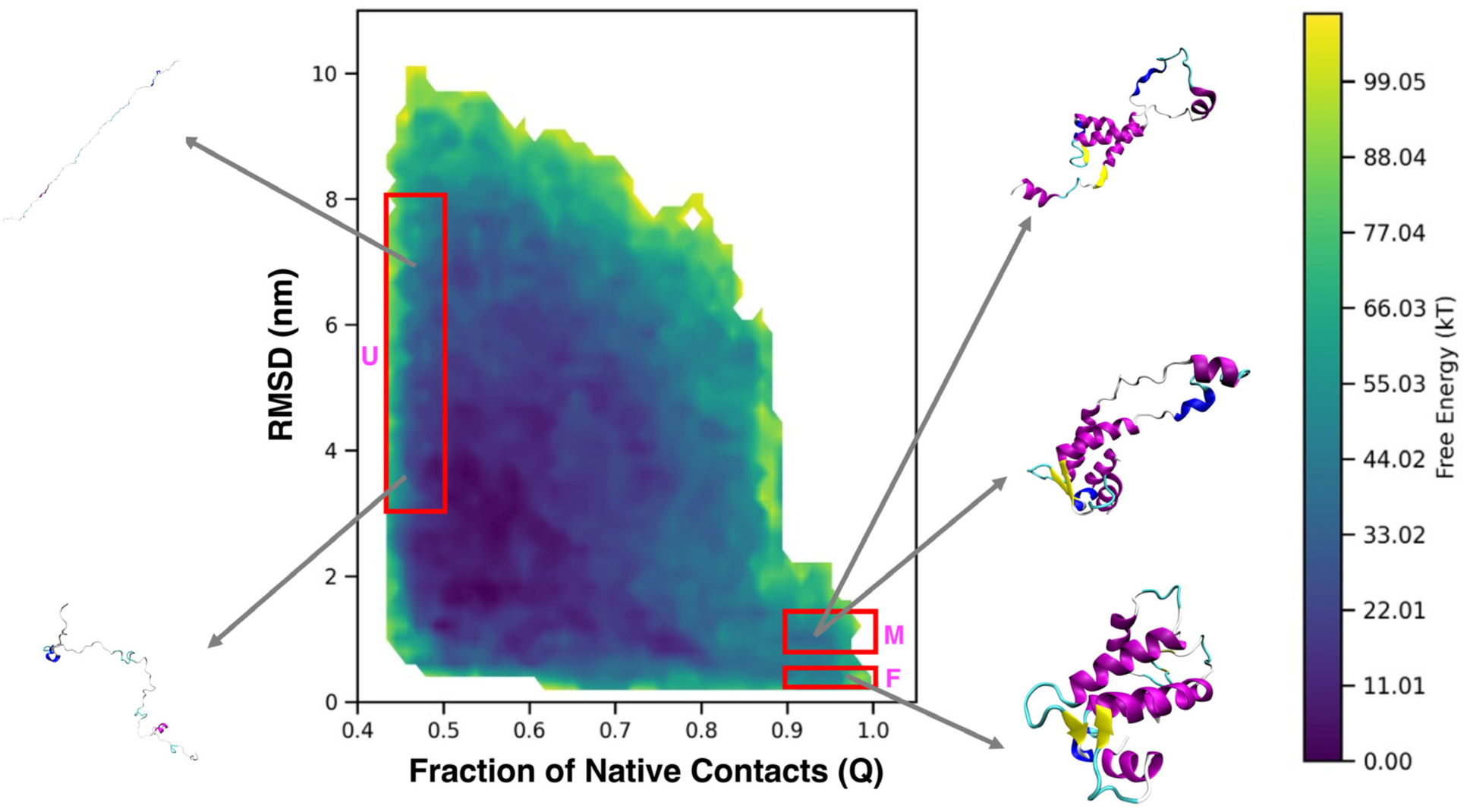
Free energy surface (FES) of Lan) at 300 K, constructed using atomistic simulations initiated from unfolded configurations. The FES is projected onto the fraction of native contacts (Q) and RMSD relative to the reference structure. Three major basins emerge: an unfolded ensemble (U, Q < 0.5, RMSD > 3 nm), a native-like folded state (F, Q > 0.9, RMSD < 0.3 nm), and a compact misfolded state (M, Q > 0.9, 0.5 < RMSD < 1.5 nm). Representative configurations sampled from each basin are included to illustrate structural diversity. These results demonstrate that spontaneous folding into native-like conformations can occur in the absence of metal ions, albeit with competition from misfolded states.

To better facilitate structural and thermodynamic analysis, we identify and define three major basins corresponding to distinct conformational ensembles: an unfolded state (U) characterized by low Q (< 0.5) and high RMSD (> 3 nm); an intermediate-to-misfolded state (M) that exhibits native-like contact formation (Q > 0.9) but deviates from the reference structure with RMSD values ranging from 0.5 to 1.5 nm; and a folded state (F) with high Q (> 0.9) and low RMSD (< 0.3 nm), corresponding to native-like conformations. Representative structural snapshots sampled from each basin (Figure 2) confirm these assignments: U-state conformations are highly disordered and largely non-helical; M-state structures retain partial or full helices in misaligned arrangements; and F-state conformations reproduce the helix–loop–helix topology observed in the crystal structure (PDB: 8DQ2).[7]

Strikingly, despite the absence of any explicit coordinating metal ions, the FES reveals a well-populated F-basin that aligns closely with the known crystal structure, demonstrating that LanM can spontaneously fold into a native-like conformation. The thermodynamic favorability and accessibility of the F-state point to a robust folding mechanism that does not require metal coordination for native structure formation. This finding strongly supports our hypothesis and demonstrates that the LanM sequence encodes sufficient energetic bias to stabilize its characteristic α-helical architecture, even under ion-free conditions. Such intrinsic foldability is particularly notable given LanM’s role as a metal-binding protein with multiple EF-hand motifs, where folding has long been presumed to be depend on ion coordination.[6, 7, 14] Moreover, this result also provides a critical baseline for our future work where we introduce ions explicitly, allowing us to dissect how lanthanide ion binding further sharpens, stabilizes, or reshapes the folding landscape. Nevertheless, in the following sections, we further dissect the nature of these states and the molecular interactions that govern transitions among them.

### Enthalpic and entropic contributions to LanM folding

To further probe the thermodynamic origins of LanM folding, we calculate temperature-dependent free energy differences (ΔF) between the unfolded (U), misfolded (M), and folded (F) states. We define the U, M, F states based on specific ranges of Q and RMSD values, as described above. Instead of calculating melting curve based on a two-state folding assumption, we report the state-to-state free energy differences ΔF_U–M_, ΔF_M–F_, and ΔF_U–F_ (where ΔF_i–j_ = ΔF_i_ – ΔF_j_) since our data suggest a more complex mechanism. **Figure 3a–c** shows these differences across temperature for each transition ΔF_U–M_, ΔF_M–F_, and ΔF_U–F_, respectively. We also extract the enthalpic (ΔH) and entropic (ΔS) components using linear fits to the van’t Hoff relation [20, 21] across the temperature range (300–450 K). This decomposition enables us to isolate the driving forces underlying each transition.

**Figure 3.**
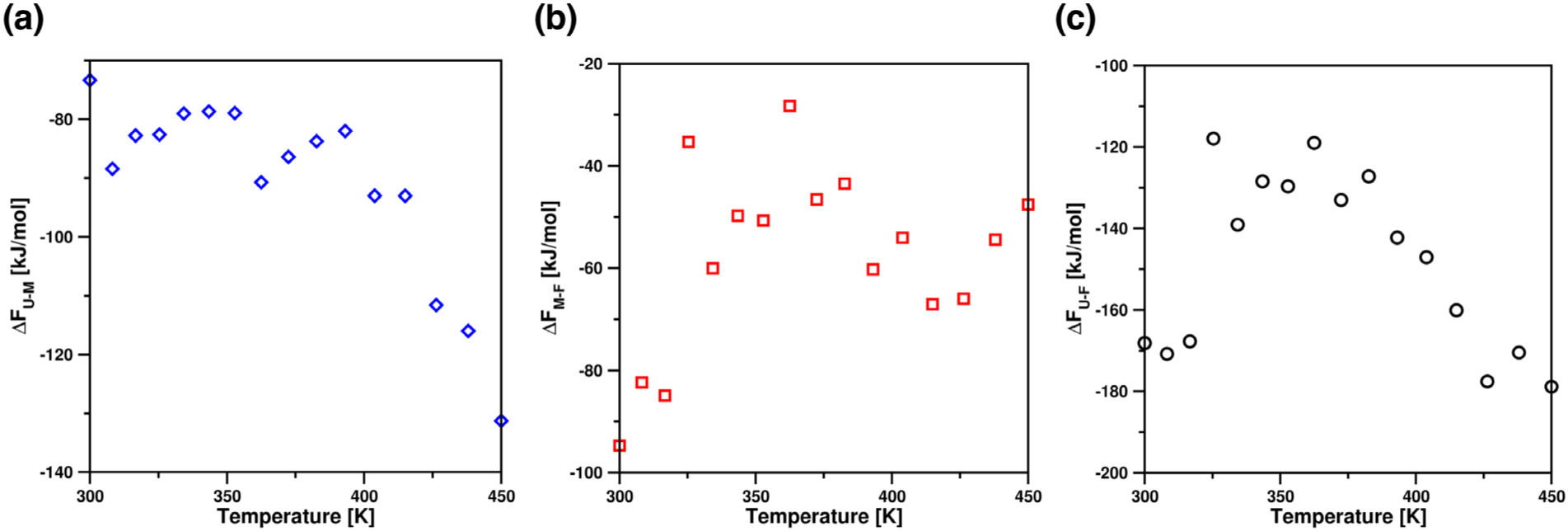
Temperature-dependent free energy differences between (a) unfolded and misfolded; (b) misfolded and folded; (c) and unfolded and folded states. For each temperature, we calculate the free energy differences using the equation ΔF_AB_ = −kT ln P_A_/P_B_, where P_A_ and P_B_ are the unbiased probabilities of finding the system in states A and B. A and B denote states U and M; M and F; and U and F for ΔF_U-M_; ΔF_M-F_; and ΔF_U-F_, respectively. The definition of the states F, M, and U in terms of the Q and RMSD variables is labeled in Figure 2.

Within the accuracy of the force field, across all transitions, ΔF decreases nonlinearly but monotonically with decreasing temperature, indicating increasing stabilization of the partially ordered states or native-like folded states at lower temperatures. The U→M transition (Figure 3a) shows a shallow ΔF profile, driven by a modest enthalpic gain (ΔH = –3.26 kJ/mol) and a favorable entropic contribution (ΔS = +0.24 kJ/mol/K), suggesting that compaction into the misfolded basin is predominantly entropically favored—consistent with the diffuse morphology of the M basin in the FES. Structurally, these misfolded states are compact but heterogeneous, with partial helices and collapsed loops that retain configurational entropy. In contrast, the M→F transition (Figure 3b) displays a steep decline in ΔF and is strongly enthalpy-driven (ΔH = –126.50 kJ/mol) while entropically penalized (ΔS = –0.19 kJ/mol/K). This indicates that the transition from misfolded intermediates to the native fold is highly specific and driven by the formation of stabilizing native contacts. The negative entropy change reflects the significant reduction in conformational freedom upon reaching the tightly ordered folded state, which appears as a narrow deep basin in the FES centered at Q > 0.9 and RMSD < 0.3 nm. The net U→F transition (Figure 3c) combines these trends, with ΔH = –129.80 kJ/mol and ΔS = +0.05 kJ/mol/K. Despite a slight entropic gain overall, the large negative enthalpy clearly dominates, pointing to enthalpy as the principal stabilizing force for the native structure. Interestingly, ΔF for both the U→F and M→F transitions show a segmented temperature dependence: values remain relatively stable between 300–315 K, exhibit a sharp increase between 315–320 K, then plateau again from 320–360 K, before decreasing at higher temperatures. This pattern may indicate temperature-dependent rearrangements in intermediate states or transitions in folding mechanisms that alter the thermodynamic balance. The M basin’s broader configuration space and the F basin’s high-contact/low-RMSD character observed in the FES align with these thermodynamic trends. Taken together, these results support a two-step folding mechanism embedded in a more complex folding landscape: an initial entropy-driven collapse into misfolded intermediates, followed by an enthalpy-dominated refinement into the native structure. This interpretation is strongly reinforced by the topography of the FES, which clearly delineates the energetic landscape associated with each state. Crucially, the fact that these transitions occur in the absence of metal ions emphasizes that the sequence alone encodes a folding pathway with near-native intermediates, although ion binding likely sharpens or accelerates this pathway.

### Structural determinants of folding across free energy basins

To uncover the structural underpinnings of the thermodynamic states identified in the FES, we analyze residue-residue contact probabilities within the unfolded (U), misfolded (M), and folded (F) basins (**Figure 4a–c**). We define the U, M, F states based on specific ranges of Q and RMSD values, as described above.

**Figure 4.**
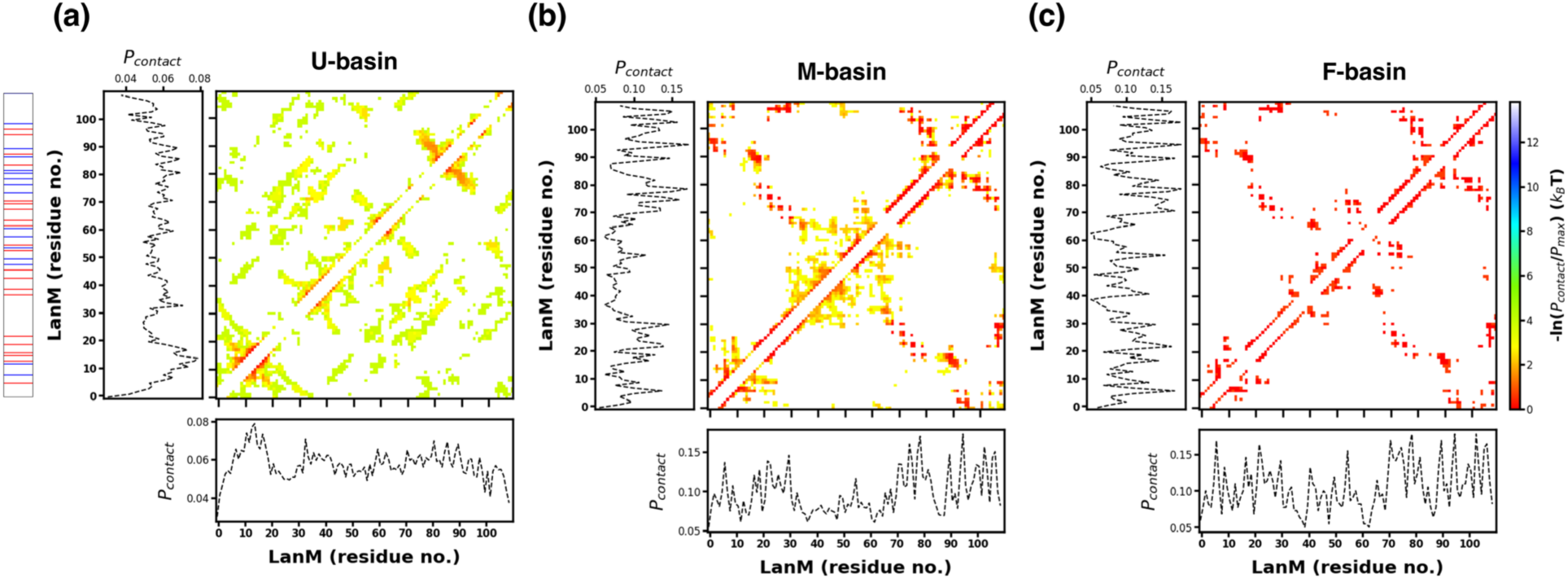
Residue–residue contact probability maps for the (a) unfolded (U), (b) misfolded (M), and (c) folded (F) basins. The definition of the states F, M, and U in terms of the Q and RMSD variables is labeled in Figure 2. Each 2D heatmap shows reweighted probability of contact between residues i and j within that basin, averaged over trajectory frames. Preferential interactions are shown in red. Contact maps reveal diffuse, heterogeneous interactions in U; partial native and non-native contacts in M, especially within individual EF-hand motifs; and high-probability native contacts across all EF-hands in F. Collectively, these patterns highlight a progressive gain in native contacts from U→M→F, supporting the thermodynamic and morphological interpretations of folding transitions.

To compute contact maps for each basin, we extract all frames from the PTWTE-WTM trajectory at 300 K whose Q and RMSD fall within the corresponding basin-defined ranges. This approach ensures consistency between thermodynamic state assignment and structural characterization. Each contact map is accompanied by contact propensities along the sequence (bottom and left panels), providing residue-level insight into folding progression. In addition, we annotate the left side of each map with the distribution of charged residues, allowing us to assess the role of electrostatic interactions in folding.

The U-basin exhibits a diffuse contact map (Figure 4a), consistent with heterogeneous, non-native interactions. Although no long-range structure is evident, several regions show weak local contacts, particularly within individual helices, suggesting the presence of transient helical motifs even in unfolded conformations. This aligns with the limited but non-zero native contact fraction (Q > 0.45) observed in the U region of the FES. Notably, contacts between oppositely charged residues appear sparse and unstructured, supporting a model of electrostatically unscreened or extended conformations. The M-basin (Figure 4b) displays a distinct pattern characterized by partially formed EF-hand motifs and several medium-range contacts, indicating partial folding and compaction. Medium-range contacts form across domains, and we observe growing contact density between regions with complementary charge, hinting at early-stage electrostatic stabilization. Although this basin retains high configurational entropy, it shows selective formation of structured motifs. This aligns with the entropy-driven thermodynamic signature of the U→M transition (Figure 3a) and the broader, shallower morphology of the M basin on the FES. The F- basin (Figure 4c), by contrast, features a sharp, highly ordered contact pattern that mirrors the crystallographic reference structure. Native contacts across all four EF-hands display high probability, with inter-helical packing evident in the off-diagonal elements. Charged residue pairings become more structured, and positively and negatively charged patches form recurrent contacts, especially in long-range regions. These observations suggest that electrostatic complementarity plays a significant role in stabilizing the final folded structure. The reorganization of tertiary contacts from M→F correlates with the strong enthalpic stabilization of folding (Figure 3b), reinforcing the cooperative and highly specific nature of native assembly.

To further dissect the physicochemical nature of folding across basins, we analyze distinct amino acid contact propensities in each state (**Figure S1**). This residue-type contact matrix complements the site-specific maps in Figure 4. Across all basins, apart from Ala residue, acidic residues (Asp, Glu) show elevated contact frequencies, likely due to LanM’s net negative charge. In the U state, these contacts are nonspecific. The M state shows broad contact distributions across polar and charged residues, reflecting heterogeneous partial folding, consistent with early-stage electrostatic collapse. In the F-basin, however, we observe enhanced contacts involving Leu, Met, and Trp – indicative of hydrophobic core formation and aromatic packing, while complementary charge interactions already established elsewhere further stabilize the folded structure, which likely together contribute to the significant enthalpic gain during the M→F transition. These trends suggest a staged folding process in which early compaction is driven by residue-type–agnostic electrostatics and later folding selects for specific polar and hydrophobic interactions.

Together, the contact analyses across FES basins reveal a progressive ordering of structural elements: from disordered ensembles in U with weak local contacts, to electrostatically collapsed partially organized EF-hand motifs in M, to a tightly packed charge-complementary native core in F. These observations reinforce the thermodynamic interpretation of folding as a two-step process – an initial entropic collapse into a metastable compact intermediate, followed by a cooperative and enthalpically driven transition into the native folded state.

### Folding pathway heterogeneity and kinetically relevant intermediates

To understand how LanM dynamically folds into its native state and whether the misfolded basin always lie on the folding pathway we analyze time-continuous folding trajectories extracted from PTWTE-WTM simulations at 300 K. We obtain these reactive trajectories by demultiplexing along the temperature ladder and identify subtrajectories that begin in the unfolded (U) basin and end in the folded (F) basin. This analysis enables us to track conformational progression along the Q–RMSD landscape and evaluate whether folding proceeds via direct or indirect routes involving the misfolded (M) basin. Overall, these trajectories offer a kinetic perspective that complements the thermodynamic and structural analyses described earlier.

In **Figure 5a**, we present the secondary structure evolution along a representative demuxed 1 μs trajectory at 300 K using DSSP assignments.[22] The simulation initiates in a largely disordered state but rapidly develops α-helical structures. As the simulation and the folding progresses, disordered segments gradually transition into more persistent helices, including coils and β- bridges, corresponding to EF-hand motifs. This stepwise emergence of helices, rather than abrupt global ordering, suggests a helix-first folding mechanism, in which local secondary structure formation precedes the establishment of long-range tertiary contacts. These trends agree with contact map observations seen in the configurations from the U and M basins (Figure 4), where local interactions nucleate early and promote subsequent compaction.

**Figure 5.**
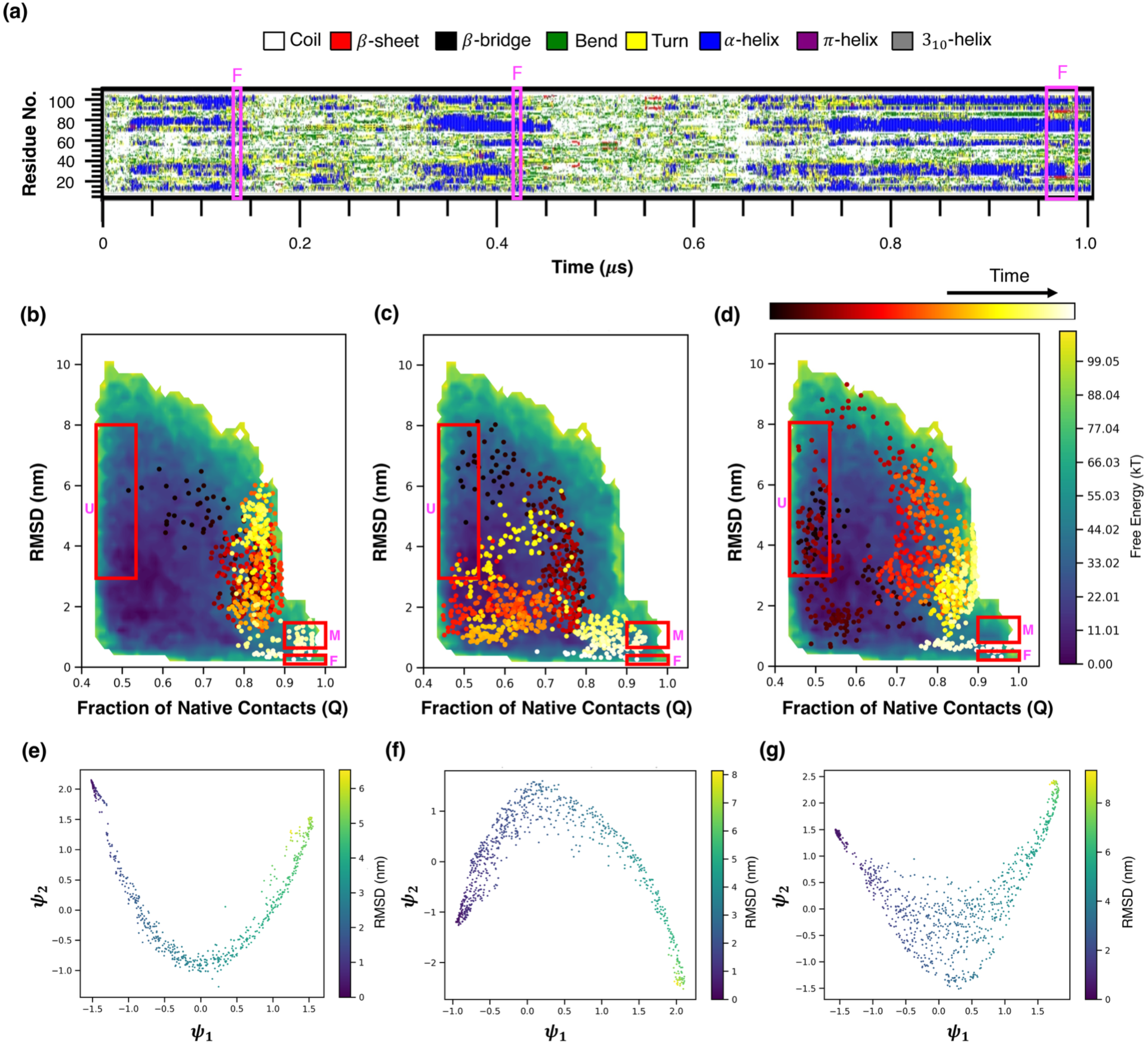
Time-resolved analysis of LanM folding reveals pathway heterogeneity and metastable intermediates. (a) Time evolution of secondary structure across a representative 1 μs folding trajectory at 300 K, as assigned by DSSP. Regions labeled “F” indicate three distinct U→F transitions. (b–d) Projections of the three reactive U→F subtrajectories (identified in panel a) onto the Q–RMSD space. Each panel corresponds to one complete folding event and illustrates distinct folding routes: panels (b) and (c) show transient sampling of the misfolded (M) basin, while panel (d) displays a direct transition from U→F. (e–g) Corresponding projections of the same trajectories onto the top two diffusion map coordinates (ψ₁, ψ₂), colored by RMSD. ψ₁ serves as a kinetically meaningful progress coordinate, differentiating unfolded (high ψ₁, high RMSD) from folded (low ψ₁, low RMSD) states. ψ₂ captures orthogonal kinetic variation and distinguishes distinct folding routes, including those passing through M and those proceeding directly. Together, these analyses reveal the presence of multiple viable pathways and support the role of M as a metastable but non-obligate intermediate in LanM folding.

Notably, the trajectory analyzed in Figure 5a undergoes three complete U→F transitions, highlighted by the labeled segments. We project these three reactive subtrajectories onto the Q– RMSD space in **Figures 5b–d**, with each panel representing a distinct folding event captured during the same simulation sampled under PTWTE-WTM conditions. In Figure 5b and 5c, the system transiently visits the misfolded (M) basin before committing to the folded state, whereas in Figure 5d, the folding path proceeds directly from U→F without sampling M state. This analysis suggests that the M basin often serve as a metastable intermediate that facilitates but is not strictly required for successful folding. Its recurrent occupation implies a kinetically accessible and potentially favorable pathway, though alternative, more direct routes are also viable.

While the Q–RMSD projections in Figures 5b–d reveal heterogeneity in folding paths and highlight the potential involvement of the misfolded (M) basin, these two biased collective variables cannot fully resolve the underlying short-time kinetic network or the structural diversity that governs LanM folding. To gain a more intrinsic view of the folding process, we apply diffusion maps to the reactive U→F subtrajectories, using RMSD-derived pairwise distances to construct a low-dimensional embedding that emphasizes slow dynamical modes. This manifold-learning approach captures the intrinsic geometry of the trajectory data, where proximity between points reflects kinetic connectivity rather than structural similarity alone. By focusing on collective transitions that evolve over long timescales, diffusion maps offer a reaction coordinate system that is both unsupervised and robust to fast fluctuations. However, we note that in this study our focus lies primarily on the structural and thermodynamic features of different folding states. Therefore, we employ diffusion maps, which provide smooth, continuous, and structurally interpretable coordinates, rather than Markov state models[23, 24] that are better suited for directly quantifying kinetic properties such as transition rates and fluxes.

In **Figures 5e–g**, we project each of these three reactive folding trajectories onto the two diffusion map coordinates (ψ₁, ψ₂), both normalized by the first leading eigenvector (ψ0), and color the points by RMSD from the crystallographic reference structure to track structural evolution. Across all three panels the resulting landscapes reveal a broadly conserved funnel-like topology and ψ₁ emerges as a global progress variable for folding: high values of ψ₁ correspond to highly disordered conformations in the unfolded (U) basin, characterized by high RMSD and low secondary structure, while low ψ₁ values localize sharply folded (F) states with compact tertiary organization and low RMSD. This monotonic trend supports ψ₁ as a reliable approximate reaction coordinate for LanM folding. In contrast, ψ₂ captures orthogonal kinetic variations and branching behavior across pathways. For instance, in Figure 5f, ψ₂ reveals a bifurcation in the trajectory’s progression: one branch veers toward the misfolded (M) basin before converging on F, while the other appears to bypass M entirely, proceeding more directly along ψ₁. This contrast becomes even clearer in Figure 5g, where the trajectory exhibits a near-linear path along ψ₁ with minimal variation along ψ₂, consistent with a direct U→F transition. These differences illustrate that the folding process can proceed through distinct routes, some involving metastable intermediates (M) and others transitioning directly from disordered to native-like states. Altogether, the diffusion map representation reconciles the heterogeneity seen in Q–RMSD projections with a kinetically coherent framework. The unfolding-to-folding transitions trace continuous paths through the ψ₁– ψ₂ space, offering a more faithful representation of pathway connectivity than any single structural descriptor. Importantly, these projections reinforce the view that LanM folding proceeds on a rugged but navigable energy landscape, where metastable intermediates are accessible and frequently visited, yet not obligatory. Folding pathways are diverse, structurally nuanced, and kinetically heterogeneous – emphasizing the importance of both local order formation and long-timescale transitions in achieving the native state.

## 3. DISCUSSION

Lanmodulin (LanM), an EF-hand protein known for its extraordinary affinity and selectivity for rare earth metal ions such as lanthanide ions, is widely presumed to fold into its native conformation only upon metal binding.[6, 7] However, our results challenge this paradigm: simulations under apo conditions reveal that LanM can spontaneously adopt a folded, native-like structure even in the complete absence of metal coordination. This finding suggests that LanM harbors an intrinsic energetic code that biases folding toward its functional topology independent of metal presence. Such behavior contrasts with classical EF-hand proteins and implies that metal binding serves more to fine-tune structure or modulate function than to initiate folding itself.

This observation has broad implications for the structural biology of metalloproteins. The emergence of persistent α-helical elements even in misfolded or unfolded ensembles, supports a helix-first folding mechanism. These local structures appear to scaffold the subsequent compaction process, promoting both structural stability and metal-readiness. As a result, LanM may exist in a preorganized, metal ion-free conformation that allows for rapid conformational response to fluctuating ion concentrations – a property potentially relevant in natural environments where metal availability is low or transient. These insights also motivate a re-examination of folding mechanisms in EF-hand proteins more broadly, especially those involved in signaling or stress responses where ion-free conformational plasticity may be functionally significant.

Methodologically, our use of PTWTE-WTM sampling enables a thermodynamic and structural dissection of the folding process. We observe a two-step mechanism: an initial entropically driven collapse into a metastable misfolded (M) basin, followed by a cooperative, enthalpically favorable reorganization into the folded (F) state. Notably, the F state forms a well-packed hydrophobic core, featuring enhanced contacts between Leu, Met, and Trp residues, alongside electrostatically complementary interactions. These structural features mirror those typically stabilized by metal ions, yet arise here in their absence, underscoring the role of side-chain encoded energetics in driving native-like topology. Further, time-resolved analyses reveal further complexity. By tracing time-resolved reactive trajectories, we uncover heterogeneous folding routes, some passing through the M basin, others bypassing it entirely. While the Q–RMSD projections capture broad basin transitions, diffusion map embeddings provide a more refined picture: ψ₁ distinguishes global folding progress, while ψ₂ resolves alternate short-time kinetic pathways. Notably, the misfolded basin appears as a distinct yet kinetically accessible branch in the manifold, suggesting its role as a metastable intermediate rather than an end state which potentially modulates the folding efficiency and responsiveness. For LanM, this framework reinforces a key insight: folding is not confined to a single reaction coordinate but is governed by a rich, multidimensional landscape shaped by both local sequence biases and global topological constraints. Our results highlight the importance of considering not only native states but also intermediate and misfolded states in understanding folding and function. In LanM, these misfolded ensembles may act as conformational reservoirs or switching intermediates in metal-sensing roles. The structural resemblance of the folded apo state to holo conformations further suggests a continuum of conformational readiness that could be exploited for function under varying environmental contexts.

These insights also bear directly on the application of LanM in rare earth element extraction and separation technologies. The finding that LanM adopts a folded-like conformation under apo conditions has important ramifications for separation system design: it implies that the protein may be pre-configured for selective metal capture, even prior to metal exposure. This structural preorganization could enable faster binding kinetics and more efficient discrimination between metal species – especially in low-concentration regimes typical of rare earth element recycling or recovery from dilute streams. Moreover, by elucidating the thermodynamic and kinetic landscape of folding, our framework lays the foundation for rational engineering of LanM variants with tailored affinities, selectivities, or responsiveness to process conditions. The rugged yet directed nature of LanM’s folding landscape highlights its versatility for deployment in complex or dynamic environments—whether immobilized on surfaces, embedded in membranes, or conjugated to scaffolds for biosensing.[12] The observed folding plasticity and pathway heterogeneity suggest robustness under diverse physicochemical constraints, expanding LanM’s utility beyond static capture agents to dynamic, reusable, and even stimuli-responsive biomolecular tools.

Taken together, our results present a nuanced picture of LanM folding that extends beyond binary models of folded/unfolded states. We highlight the value of trajectory-based analyses in uncovering folding intermediates, challenge long-standing assumptions about the role of cofactors in metalloprotein structure, and provide a generalizable methodology for studying flexible, conditionally disordered systems. We anticipate that this approach will be useful for probing the folding of other cofactor-regulated proteins and for understanding how structure, function, and environmental sensing are integrated in dynamic biological systems.

## 4. METHODS

### Model and Simulation details

Following the methodology of Zerze et al.,[25] we use parallel tempering in the well-tempered ensemble combined with well-tempered metadynamics (PTWTE-WTM)[26–31] to sample the folding landscape of the lanmodulin (LanM) protein. We perform all PTWTE-WTM simulations using GROMACS (version 2022.5)[32, 33] patched with the PLUMED (version 2.9.0) enhanced sampling plugin[34, 35] for metadynamics sampling. We employ 16 temperature replicas, spaced geometrically between 300 K and 450 K, with replica swap attempts every 1000 steps.

In the PTWTE framework, we bias the system’s potential energy using Gaussian kernels with an initial width of 500 kJ/mol and a height of 1.0 kJ/mol. Gaussians are deposited every 2000 steps, with a bias factor of 16.

For the WTM component, we use root-mean-square deviation (RMSD) and a similarity-based order parameter Q as collective variables. We intentionally exclude ion–protein interactions and do not include metal coordination distances as collective variables, because we aim to probe the intrinsic thermodynamic tendency of LanM to fold in the absence of lanthanide ions. By biasing the simulations along RMSD and Q – two structural descriptors that capture native-state formation – we directly examine the protein’s inherent folding landscape. This approach allows us to test whether LanM adopts its folded conformation without relying on metal binding.

We define both relative to the folded structure of LanM, selected from the X-ray crystal structure of *Hansschlegelia quercus* LanM (PDB ID: 8DQ2).[7] RMSD includes only C_α_ atoms, and Q captures atomic contacts within native secondary structure elements. The order parameter Q is defined as

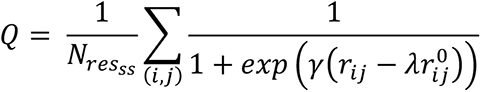

by adapting the generalized definition of Q.[36] The sum runs over *N_res_ss__*, which includes all heavy-atom pairs (*i,j*) in the secondary structure (excluding intra-residue contacts) that are within 0.5 nm in the reference native structure; which reinforces the role of backbone hydrogen bonding and local helix-helix contacts in stabilizing the folded state. 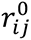 and 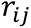 are the distances between *i* and *j* in the reference native structure and in any given instantaneous configuration, respectively. The smoothing function *γ* is taken as 50 nm^-1^ and the adjustable parameter *λ* is taken as 1.5.[36]

We apply bias to RMSD and Q using Gaussian kernels with widths of 0.05 and 0.01, respectively, and a height of 1.8 kJ/mol. Gaussians are deposited every 500 steps with a bias factor of 35. To avoid artifacts near the bounds of Q, we restrain it between 0.01 and 0.99 using harmonic walls with a spring constant of 10,000 kJ/mol and also set the metadynamics force on Q to zero outside this interval.[37, 38]

To construct unfolded starting structures, we run coarse-grained parallel tempering simulations[26] in LAMMPS (Oct. 2020 version)[39] using the HPS-Urry protein model[40] – a coarse-grained protein model that we have successfully used in our previous studies to investigate sequence-dependent properties of intrinsically disordered proteins.[41–43] We simulate geometrically spaced 16 temperature replicas from 300 K to 450 K for 0.5 μs each, using Langevin dynamics[44] with a 10 fs timestep, 1000 time steps damping parameter and replica swap attempts every 100 steps. We discard the first 0.25 μs of each trajectory as equilibration. Based on radius of gyration analysis (**Figure S2**), we extract a single representative unfolded structure and back-map it to an atomistic model using MODELLER.[45, 46]

Next, we solvate this single copy configuration in a 9 nm cubic box, add Na⁺ ions for neutrality, and set the NaCl concentration to 100 mM. We use the AMBER03ws force field[47] for LanM, TIP4P/2005[48] for water, and improved ion parameters from Lou and Roux.[49] We subject this initial configuration, with periodic boundary conditions in x, y, and z directions, to steepest descent energy minimization to remove overlaps. Following minimization, we equilibrate the system with unbiased 1 ns NVT simulation (T = 300 K) followed by unbiased 2 ns NPT simulation (T = 300 K, P = 1 bar). We use the final NPT snapshot as the starting point for PTWTE-WTM simulations.

We perform 1 μs of PTWTE-WTM production per replica under NPT conditions. We maintain the respective replica temperature using the stochastic velocity-rescale thermostat (τ = 0.4 ps),[50] and pressure at 1 bar using the Parrinello–Rahman barostat (τ = 2 ps).[51] We treat electrostatics with particle mesh Ewald (PME)[52] and truncate both electrostatics and Lennard-Jones[53] interactions at 1 nm, applying analytical long-range corrections to the latter.[54] Although we constrain bonds involving hydrogens with LINCS,[55] we integrate the equations of motion using a 1 fs timestep and leapfrog algorithm. We discard the first 0.1 μs per replica as equilibration and store frames every 200 ps, resulting in 4500 configurations per replica for analysis.

### Convergence

As a rigorous test of convergence, we perform two independent PTWTE-WTM simulations of the LanM protein. In the first, we initialize all replicas from a fully unfolded configuration (as described above), while in the second, we start all replicas from the correctly folded structure, selected from the 8DQ2 PDB entry. For both starting conditions, **Figure S3** shows the time evolution of RMSD and Q at the 300 K replica. Evidently, the initial configurations undergo several round trips through the entire RMSD and Q space, which indicates efficient conformational exchange between replicas. We also use the simulation that starts from the folded configuration to quantify deviations between the two cases. **Figure S4** presents the free energy surfaces as a function of RMSD and Q, along with their difference, which we interpret as an estimate of error. The resulting free energy surfaces, projected onto the Q–RMSD space, show strong agreement in the folded basin (high Q, low RMSD), which represents the thermodynamically most relevant region. We observe that the differences between the two simulations generally remain within ±10*kT* across the central folding landscape, indicating consistent sampling of intermediate states. Larger discrepancies appear only in the peripheral regions of high RMSD and low Q, which correspond to entropically broad and sparsely visited unfolded conformations. These variations are expected due to the high-dimensional conformational space and the stochastic nature of bias deposition in protein folding simulations. Given the agreement across folded and intermediate regions, and the use of bidirectional sampling from both folded and unfolded initial states, we consider the simulations to be near-converged in the thermodynamically relevant portions of the folding landscape. In the main text, we report results based on the simulation that starts from the unfolded configuration.

### Analyses

We quantify the free energy surface (FES) of the LanM protein using two collective variables, Q and RMSD, both of which we bias during the PTWTE-WTM simulations. To obtain the unbiased two-dimensional probability density P(Q,RMSD), we apply the reweighting scheme introduced by Tiwary and Parrinello,[56] which accounts for all deposited biases on Q and RMSD coordinates. We then compute the free energy as:

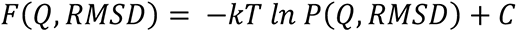

where kT is the thermal energy (the product of the Boltzmann constant and temperature), and C is an arbitrary additive constant.

To gain insights into the folding pathways, we analyze reactive trajectories by computing Q and RMSD along continuous trajectories—i.e., trajectories that exchange temperature but remain continuous in time. We generate these time-continuous trajectories by demultiplexing the parallel-tempered simulations using the demux code in the GROMACS package. Within these demultiplexed trajectories, we: (a) track the evolution of secondary structure using the DSSP library[22] via the gmx do_dssp utility in GROMACS package, (b) identify subtrajectories that begin in the unfolded configuration and end in the correctly folded conformation, and (c) analyze how intramolecular contacts form as the system transitions from unfolded to folded state. For contact analysis, we define two residues *i* and *j* (with sequence separation ∣*i*−*j*∣>3) as being in contact if any of their heavy atoms lie within 0.6 nm of each other. We select this cutoff based as it captures a range of interaction types, including hydrogen bonds (<0.35 nm), van der Waals contacts, and salt bridges (<0.6 nm).[57]

To complement our structural analysis, we compute free energy differences between distinct conformational basins across a range of temperatures. This state-resolved ΔF analysis allows us to capture the underlying thermodynamics without relying on a simplified two-state assumption. By evaluating how these free energy differences vary with temperature, we estimate the corresponding enthalpic and entropic contributions. This decomposition provides key insight into the thermodynamic driving forces underlying conformational transitions and how temperature modulates the relative stability of various regions of the energy landscape.

To further explore the dynamical organization of the folding process, we also apply diffusion map analysis[58] to the ensemble of reactive subtrajectories to uncover low-dimensional collective variables that capture the dominant conformational transitions during folding. Following the methodology of Alvarado et al.,[59] we construct a diffusion kernel using a Gaussian weighting function applied to pairwise structural distances, where distance is computed using RMSD over C_α_ atoms between translationally and rotationally aligned LanM coordinates in frames *i* and *j* of the trajectory. We select the kernel bandwidth *∈* as 3.0 and diffusion map exponent *α* as 0.3 based on the median of the distance distribution to balance locality and global structure.[60] We compute the top three eigenvectors of the transition probability matrix and normalize the first two nontrivial diffusion components by the leading eigenvector, which corresponds to the stationary distribution. We then use the resulting scaled coordinates to construct the diffusion map projections, which together account for the majority of short-time kinetic variance in the dataset. These coordinates serve as effective reaction coordinates that reveal dominant folding pathways, intermediate metastable states, and conformational bottlenecks. This framework enables us to move beyond traditional order parameters and visualize the folding landscape in a manner that naturally clusters kinetically similar configurational microstates in close proximity. We note that we perform this diffusion map analysis by adopting the DMAPS library[61] with in-house python scripts.

## AUTHOR CONTRIBUTIONS

Utkarsh Kapoor: Conceptualization; investigation; funding acquisition; writing – original draft, review and editing; formal analysis; visualization.

## AUTHOR INFORMATION

*Corresponding Author* Utkarsh Kapoor – Department of Chemical and Biomedical Engineering, University of Wyoming, Laramie WY 82071, United States;

## DATA AVAILABILITY

Raw data and code are available upon reasonable request.

## SUPPORTING INFORMATION

Supporting figures showing contact analysis based on residue-type, the sampling of metadynamics collective variables as a function of time and the difference in the free energies have been included.

## ACKNOWLEDGMENTS

We extend our sincere gratitude to Dr. Karen E. Wawrousek and Dr. Caleb M. Hill for their insightful discussions. We also acknowledge the financial support provided by the University of Wyoming, particularly through the Engineering Initiative and Research Excellence seed grants. This research was partially completed with advanced computing resources generously provided by the University of Wyoming Advanced Research Computing Center (UW-ARCC), specifically the MedicineBow supercomputing cluster. Furthermore, we recognize the computing time on the Derecho system (doi:10.5065/qx9a-pg09) supported by the NSF National Center for Atmospheric Research (NCAR) at the NSF NCAR-Wyoming Supercomputing Center, sponsored by the National Science Foundation and the State of Wyoming.

