## Supplementary material for "Folding Thermodynamics and Pathway Heterogeneity of Lanmodulin from Atomistic Simulations": SI

***Utkarsh Kapoor*[1*]**

*1. Department of Chemical and Biomedical Engineering, University of Wyoming, Laramie, WY 82071, United States*

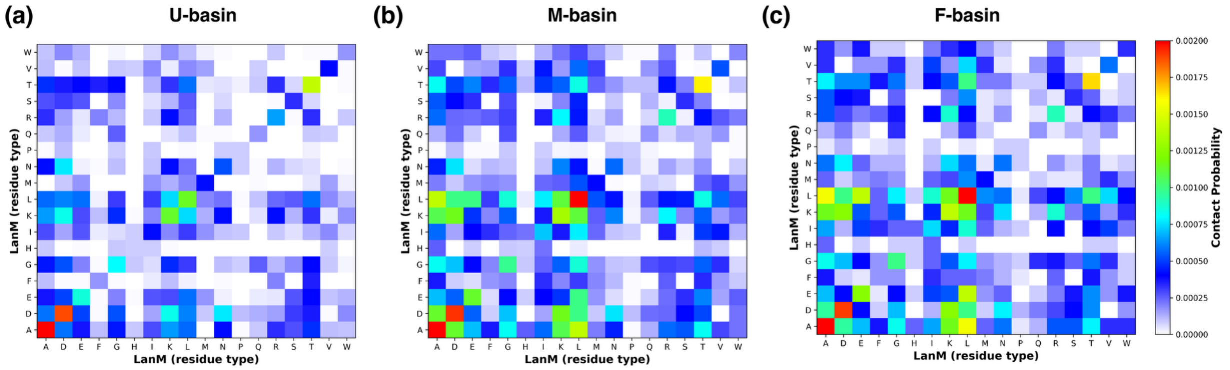


***Figure S1.*** *Residue-type based contact frequency maps for LanM in the (a) unfolded (U), (b) misfolded (M), and (c) folded (F) basins, computed from trajectory frames assigned to each basin. Preferential interactions are shown in red.*

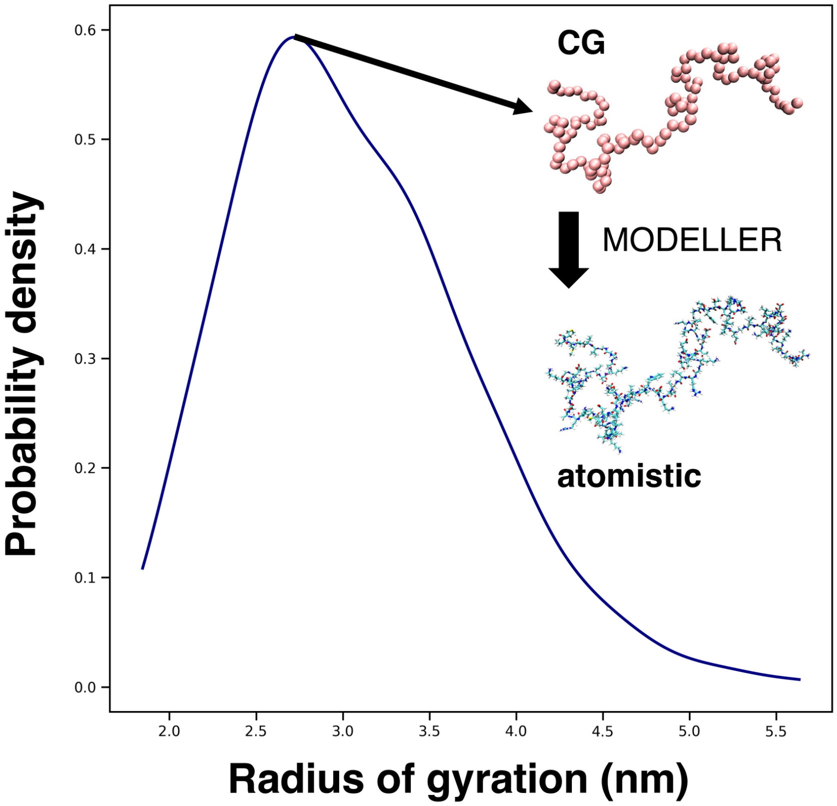


***Figure S2.*** *Distribution of radius of gyration values at 300 K sampled from coarse-grained parallel tempering simulations of LanM using the HPS-Urry model. A representative structure from the most probable region of the distribution was extracted and back-mapped to atomistic resolution using MODELLER, enabling construction of native-like unfolded conformations for atomistic PTWTE-WTM simulations. This workflow allows generation of physically relevant unfolded starting structure of LanM without artificial bias.*

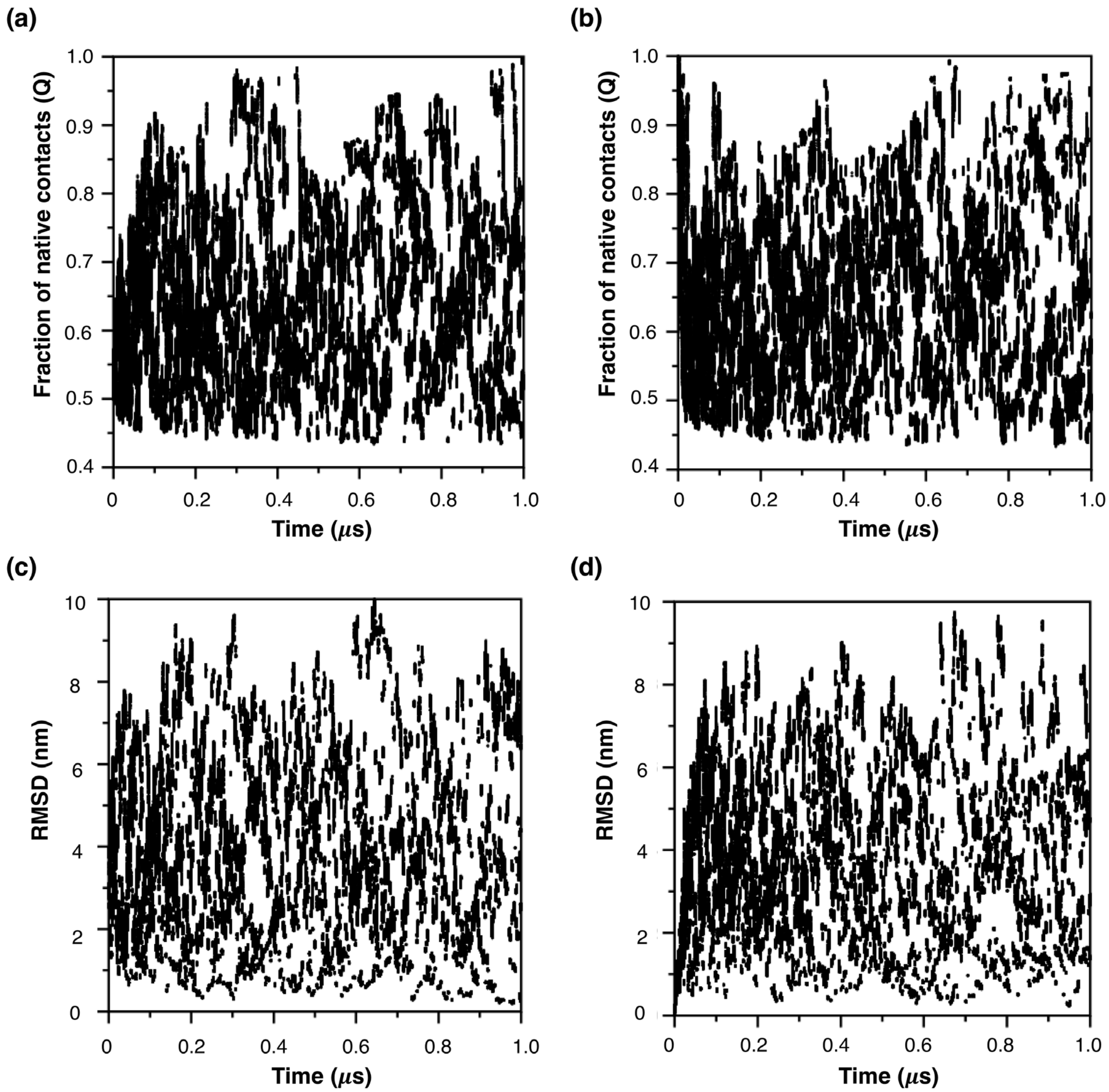


***Figure S3.*** *Sampling of Q (top panel), and RMSD (bottom panel) as a function of time for LanM starting from two independent initial conditions at 300 K (left: unfolded configuration, right: reference native configuration).*

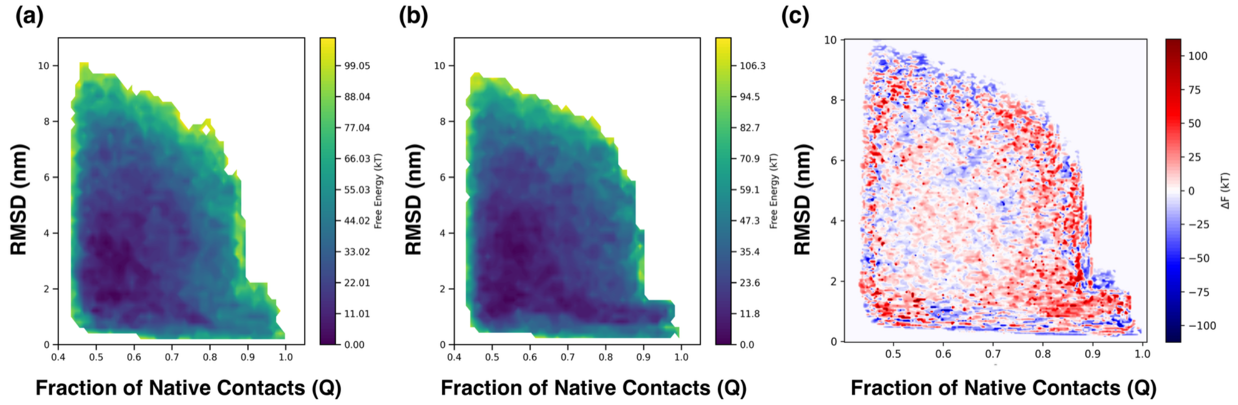


***Figure S4.*** *FES of LanM as a function of Q and RMSD for different initial conditions at 300 K (a: starting from unfolded configuration and b: starting from reference native configuration). The absolute deviation between the FES is reported as error in c.*
